# Individual neuronal subtypes control initial myelin sheath growth and stabilization

**DOI:** 10.1101/809996

**Authors:** Heather N. Nelson, Anthony J. Treichel, Erin N. Eggum, Madeline R. Martell, Amanda J. Kaiser, Allie G. Trudel, James R. Gronseth, Samantha T. Maas, Silas Bergen, Jacob H. Hines

## Abstract

**Background:** In the developing central nervous system, pre-myelinating oligodendrocytes sample candidate nerve axons by extending and retracting process extensions. Some contacts stabilize, leading to the initiation of axon wrapping, nascent myelin sheath formation, concentric wrapping and sheath elongation, and sheath stabilization or pruning by oligodendrocytes. Although axonal signals influence the overall process of myelination, the precise oligodendrocyte behaviors that require signaling from axons are not completely understood. In this study, we investigated whether oligodendrocyte behaviors during the early events of myelination are mediated by an oligodendrocyte-intrinsic myelination program or are over-ridden by axonal factors.

**Methods:** To address this, we utilized in vivo time-lapse imaging in embryonic and larval zebrafish spinal cord during the initial hours and days of axon wrapping and myelination. Transgenic reporter lines marked individual axon subtypes or oligodendrocyte membranes.

**Results:** In the larval zebrafish spinal cord, individual axon subtypes supported distinct nascent sheath growth rates and stabilization frequencies. Oligodendrocytes ensheathed individual axon subtypes at different rates during a two-day period after initial axon wrapping. When descending reticulospinal axons were ablated, local spinal axons supported a constant ensheathment rate despite the increased ratio of oligodendrocytes to target axons.

**Conclusion:** We conclude that properties of individual axon subtypes instruct oligodendrocyte behaviors during initial stages of myelination by differentially controlling nascent sheath growth and stabilization.

## Background

Ensheathment of nerve axons with myelin is an essential process during vertebrate neural development. Oligodendrocytes are specialized central nervous system (CNS) ensheathing cells that extend multiple membrane processes to sample candidate axons, initiate and perform spiral axon wrapping, then extend along axons to form a mature myelin sheath (reviewed by [1,2]). Alternatively, oligodendrocytes can retract and prune myelin sheaths [3–5], as well as modify the thickness of individual sheaths in response to external stimuli [6]. These complex steps take place over the course of days or weeks, and ultimately are responsible for mature myelin sheaths being positioned at the right location, time, and parameters [7–9]. Numerous distinct cell behaviors and specialized mechanisms are deployed during these steps, including dynamic process extension and retraction, cell-type recognition, cell adhesion, and protrusive myelin membrane growth for concentric wrapping and extension along axons (reviewed by [10–12]). No single axonal factor has been identified that is required for myelination, but many regulate the abundance, length, and thickness of myelin sheaths [9,12–15]. How and when these factors regulate stage-specific oligodendrocyte cell behaviors, leading to the complex myelination patterns observed in the CNS, is not well understood.

The discoveries that cultured oligodendrocytes form immature myelin onto chemically fixed axons and synthetic fibrils indicate that oligodendrocytes possess an intrinsic ensheathment program that is capable of executing at least some of the cell behaviors and stages of myelination ([16–19]. The extent to which oligodendrocytes utilize axon-derived cues in vivo is less clear, but it is possible that the first steps and behaviors mediating sheath formation and initial stabilization could occur without molecular instruction from axons. In vitro, oligodendrocytes require synthetic fibrils to possess a minimal diameter of 0.3 μm in order to initiate sheath formation [17]. Regulation of axon diameter by radial growth (or lack thereof) could therefore define axons as either permissive or non-permissive for myelination, after which continued wrapping, stabilization, and sheath maturation could be supported by axon-derived molecular cues. In this way, an oligodendrocyte-intrinsic ensheathment program could mediate initial sheath formation based on the permissive diameter of an axon, followed by axon-derived signals that instruct later oligodendrocyte behaviors to complete and refine the myelination process [20]. In support of this model, an artificial increase in axon diameter by genetic deletion of PTEN is sufficient to induce oligodendrocytes to myelinate cerebellar granule cell axons, which are normally unmyelinated [21]. Furthermore, a recent study demonstrated that oligodendrocytes continue to initiate myelination in the optic nerve in the absence of dynamic neuronal signaling [22]. Numerous studies collectively support the notion that oligodendrocytes are intrinsically programmed to wrap any axon of suitable caliber, but the extent to which these mechanisms facilitate the unique spatial and temporal profiles of CNS myelination in vivo is less clear, and may be region- or axon subtype-specific.

The CNS comprises many distinct neuronal subtypes, each possessing specific genetic profiles, structural morphologies, and functional connectivity. Oligodendrocytes stereotypically myelinate some axon subtypes but not others, however the cellular interactions forming this heterogenous myelin landscape are unknown. In this study we provide evidence that the earliest steps of sheath formation, stabilization, and growth are influenced by properties of individual axons. Using in vivo imaging in the larval zebrafish spinal cord, we found that distinct axon subtypes differentially controlled the rate of nascent sheath growth and initial stabilization. Consequently, the myelin ensheathment of distinct axon subtypes proceeded at different rates during the 48 hours following initial wrapping. When a subset of target axons was ablated, remaining axon subtypes maintained consistent ensheathment rates despite the increased ratio of oligodendrocytes to myelin-competent axons. Taken together, these findings support the hypothesis that properties of individual axons are sufficient to influence or override the oligodendrocyte-intrinsic ensheathment program by regulating initial sheath growth and stabilization at the onset of myelination in vivo.

## Results

To determine how oligodendrocyte behaviors are influenced by different axon subtypes, we first identified myelin-competent target axons within the larval zebrafish spinal cord. Previous studies established reticulospinal neurons and commissural primary ascending (CoPA) interneurons as myelinated axon subtypes [5,23]. In the spinal cords of 3-5 dpf larvae, we observed *Tg(tbx16:EGFP)* reporter expression restricted to CoPA neurons [24]. CoPA axons projected ipsilaterally toward the ventral floor plate, crossed the ventral midline, and extended contralaterally and anteriorly into and within the dorsal longitudinal fasciculus (DLF) (Additional Fig. 1a). When outcrossed to *Tg(sox10:RFP-CaaX)*, which marks oligodendrocyte processes with membrane-tethered RFP, we observed ensheathment of EGFP^+^ CoPA distal axon segments located within the DLF (Fig. 1a). Oligodendrocytes did not myelinate ipsilateral axon segments, but myelinated all contralateral axon segments. We also utilized the *Tg(isl1[ss]:Gal4, UAS:DsRed)* reporter line to label Rohon-Beard (RB) neurons, a second population of local spinal neuron axons. RB sensory neuron axons also extended within the DLF, adjacent but immediately dorsal to the position of CoPA axons. When outcrossed to *Tg(nkx2.2a:EGFP-CaaX)*, which marks oligodendrocyte processes with membrane-tethered EGFP, we observed ensheathment of DsRed+ RB axons (Fig. 1a) [23,25].

**Fig. 1.**
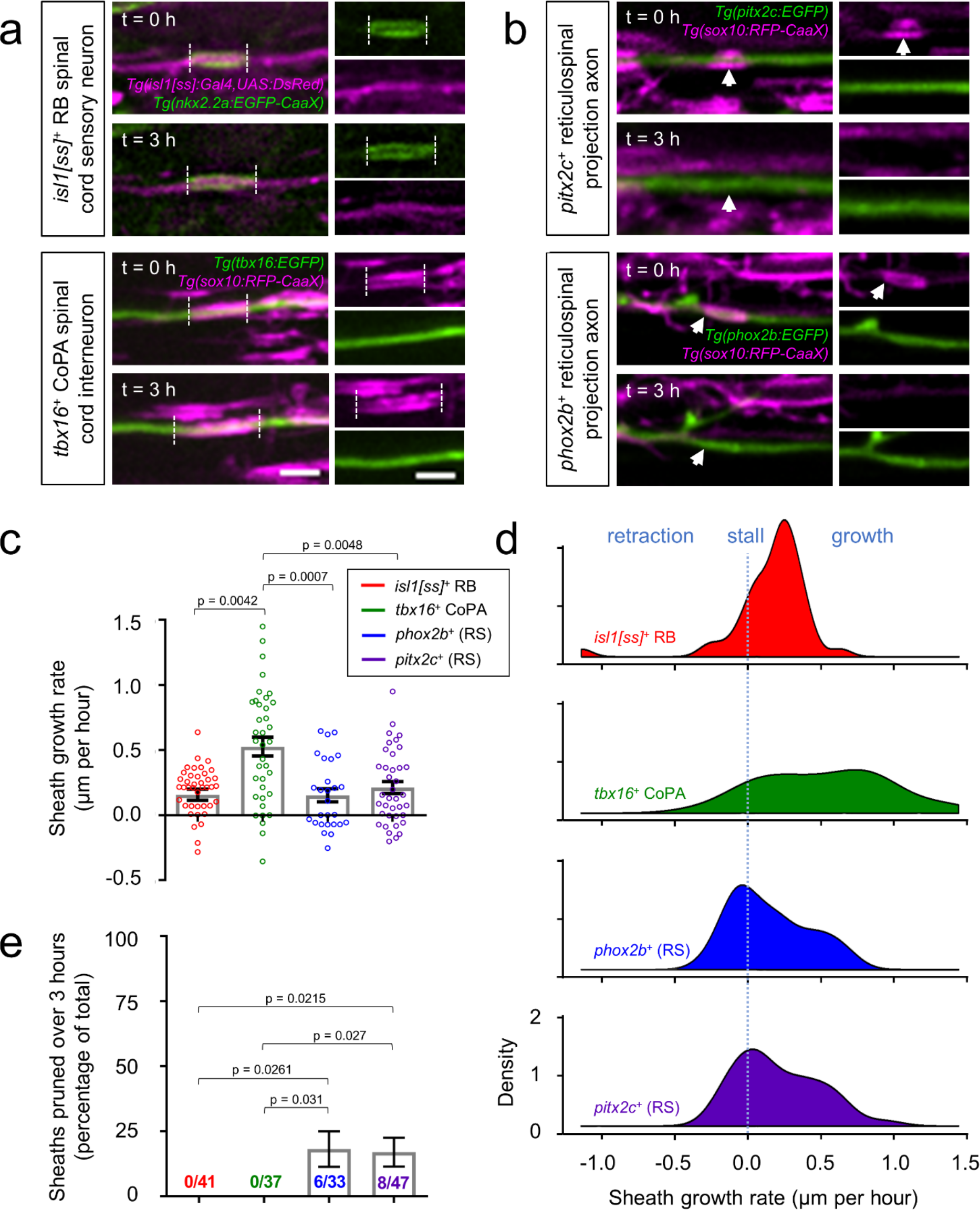
Individual sheath extension rates and pruning frequencies are axon subtype-specific. **a-b** Lateral view images of the larval spinal cord show sheath extension **(a)** and sheath pruning **(b)** during a 3-hour time course. Transgenic reporter expression (indicated) marks individual axons or oligodendrocyte membranes. Dashed lines illustrate individual sheath lengths and arrowheads denote sheath pruning events. Images are confocal maximum z-projections with dorsal up and anterior left. Scale bar = 3 μm. **c-d** Plots show mean growth rates (**c**) and distributions of individual sheath growth rates (**d**). For (**c**), error bars represent the mean ± SEM. For **(d)**, the y-axis approximates the density of data points that exhibited the corresponding growth rate. Larger y-values represent a larger proportion of data points exhibiting this growth rate. n (# of sheaths) = 27 (*phox2b*^+^), 37 (*tbx16*^+^), 38 (*pitx2c*^+^), 41 (*isl1[ss]*+). **e** Quantification of the percentage of total sheaths that pruned during a 3-hour time course grouped by axon subtype. n (sheaths) = 33 (*phox2b*^+^), 37 (*tbx16*^+^), 47 (*pitx2c*^+^), 41 (*isl1[ss]*+). Overall chi-square test of difference in percent pruning = 0.0018. Pairwise p-values were generated using Dunn’s multiple comparisons (**c**) or using Tukey-adjusted pairwise comparisons (**e**).

Spinal cord oligodendrocytes myelinate axons projecting from both local spinal cord neurons and descending reticulospinal neuron populations. We previously identified *phox2b*^+^ neurons as hindbrain reticulospinal axons that become myelinated within and along the length of the ventral spinal cord medial longitudinal fasciculus (Fig. 1b, Additional Fig. 1c) [5]. To identify additional reticulospinal myelinated axon subtypes, we crossed *Tg(sox10:RFP-CaaX)* to the *Tg(pitx2c:EGFP)* reporter line, which marks the rostral-most reticulospinal neurons of the nucleus of the medial longitudinal fasciculus (nMLF) [26]. nMLF cell bodies, resident to the ventral midbrain, projected axons into and posteriorly across the length of the ventral spinal cord (Additional Fig. 1d), consistent with an established role for the nMLF in coordinating locomotor responses to visual input [27–29]. In 3-5 dpf larvae, we observed *sox10*^+^ sheaths wrapping EGFP^+^ nMLF axons located in the ventral spinal cord (Fig. 1b).

### Subtype-specific rates of initial ensheathment and sheath pruning

If the early events of myelination are mediated by an oligodendrocyte-intrinsic ensheathment program, we predicted that the behavior of newly formed (nascent) individual myelin sheaths would be independent of the axon subtype ensheathed. In contrast, if unique properties of individual axons play an instructive role during these early stages, we hypothesized that initial sheath growth and stabilization would differ depending on the axon subtype being wrapped. To distinguish between these possibilities, we identified nascent sheaths wrapping transgene-labeled axons and tracked progression over three hours (Fig. 1). We defined nascent sheaths as those less than 5 μm in length. This criterion was based on the previous finding that sheaths that are initially stabilized extend to greater than 5 μm within three hours or less [5]. Therefore, in our study, observed sheaths less than 5 μm in length had most likely formed within the previous three hours. Nascent sheaths wrapping *isl1[ss]*^+^ RB sensory axons extended at an average rate of 0.2 μm/h (Fig. 1a,c). In comparison, other sheaths within the DLF that wrapped *tbx16*^+^ CoPA axons extended at a comparatively increased average rate of 0.5 μm/h (Fig. 1a,c). Nascent sheaths wrapping both *pitx2c*^+^ and *phox2b*^+^ ventral-residing reticulospinal axons extended at an average rate of 0.2 μm/h (Fig. 1c). In addition to subtype-specific differences in the average sheath growth rate, we also noted subtype-specific differences in the distribution of the individual sheath growth rates that composed the average (Fig. 1d). Within the dorsal tract, *isl1[ss]*^+^ RB sensory axons predominantly exhibited slow sheath growth (mostly 0.2 μm/h or less), whereas *tbx16*^+^ CoPA axons had the ability to support more rapid sheath growth (up to 1.4 μm/h). Within the ventral tract, sheaths wrapping both *pitx2c*^+^ and *phox2b*^+^ reticulospinal axons exhibited a tendency to stall or retract, however sheaths that grew largely did so at a moderate rate (~0.4 μm/h). Therefore, although both dorsal sensory and ventral reticulospinal axons had comparable average sheath growth rates (0.2 μm/h), sheaths wrapping dorsal sensory axons achieved this through slow growth, whereas sheaths wrapping ventral reticulospinal axons performed a combination of retractions, stalls, and moderate growth to achieve the same average growth rate. Together, these data reveal axon subtype-specific differences in overall sheath growth rate and further imply that individual axon subtypes instruct different patterns of sheath growth.

Previous studies indicate that some initial axon wrapping attempts can undergo retraction and pruning, motivating us to ask whether the outcome of sheath stabilization versus pruning could be axon subtype-dependent [3–5]. During the three-hour imaging window, oligodendrocytes stabilized 100% of sheaths wrapping *tbx16*^+^ CoPA axons (37/37) and *isl1[ss] +* RB sensory axons (41/41) (Fig. 1a,e). In contrast, 18% (6/33) and 17% (8/47) of nascent sheaths wrapping *phox2b*^+^ and *pitx2c*^+^ reticulospinal axons were pruned during the imaging period, respectively (Fig. 1b,e).

Do differential nascent sheath growth rates and pruning frequencies result in axon subtype-specific differences in overall ensheathment rate? We next tracked the initial 48-hour progression of axon ensheathment on transgene-labeled subtypes. For each, we identified a baseline time point (t = 0) based on the earliest appearance of nascent sheaths within a defined spinal cord segment, and subsequently acquired images within the same spinal segment 24 and 48 hours later. Oligodendrocytes did not increase the overall percent ensheathment of *isl1[ss]*^+^ RB sensory axons over the 48-hour period post initial wrapping (Fig. 2a,c). In stark contrast, oligodendrocytes rapidly myelinated *tbx16*^+^ CoPA axons and arrived at near complete ensheathment (92.0 ± 1.2% of total axon length) during the same 48-hour period post initial wrapping (Fig. 2b,c). In comparison to the near full ensheathment of *tbx16*^+^ CoPA axons observed 48 hours post initial wrapping, oligodendrocytes ensheathed 43.8 ± 4.8% and 32.5 ± 8.7% of the length of *pitx2c*^+^ and *phox2+* spinal projection axons, respectively (Fig. 2c). Taken together, these results indicate that properties of individual axon subtypes control the initial rate of overall ensheathment by differentially regulating both individual sheath growth and stabilization.

**Fig. 2.**
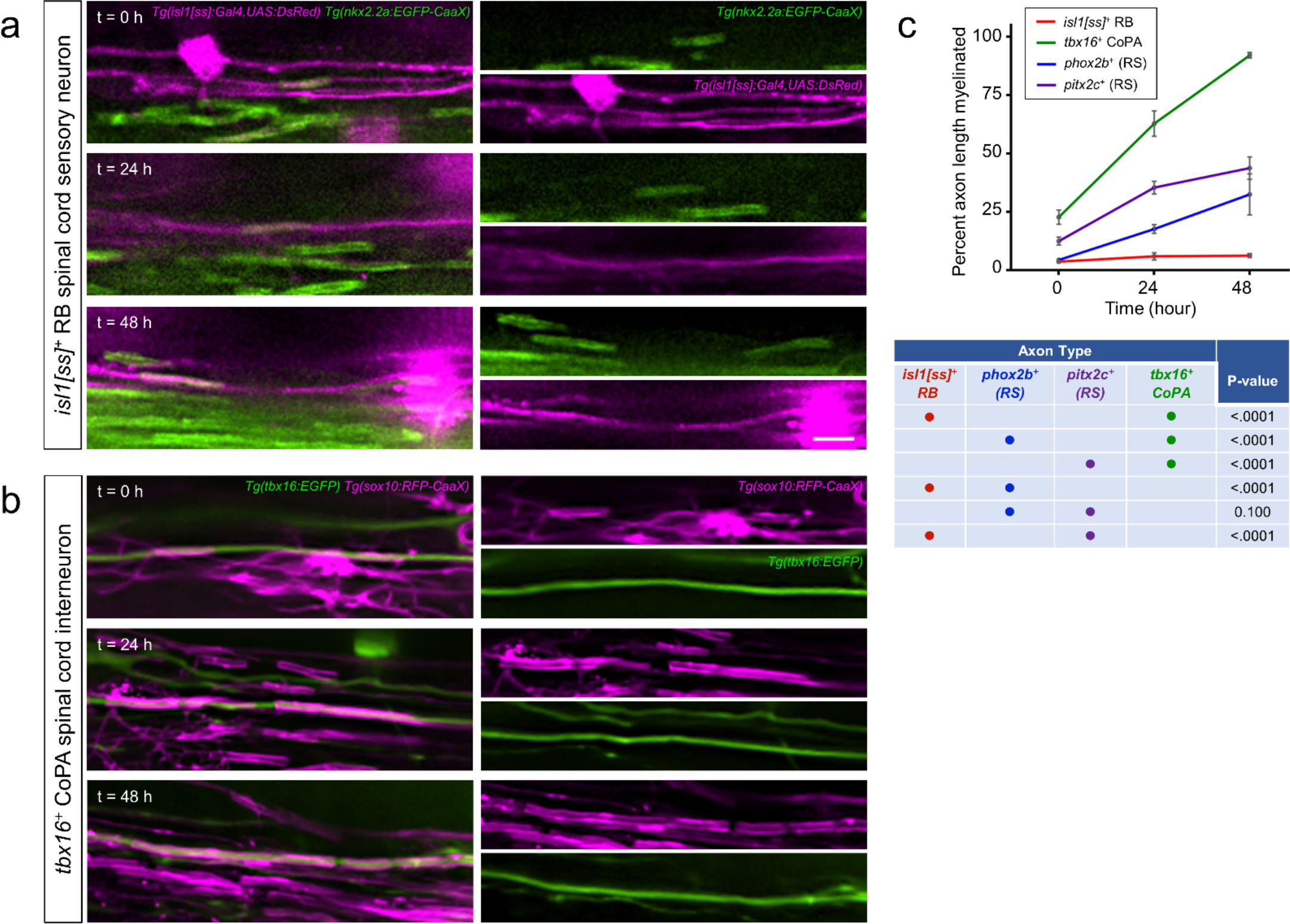
Individual axon subtypes support differential initial ensheathment rates. **a-b** Lateral view images show the progression of myelination on individual spinal axons during the 48-hour period following initial wrapping. Images were acquired within a constant spinal cord segment at each time point. Note that although RB and CoPA spinal axons are both located within the spinal cord DLF, RB sensory axon myelination remained sparse and incomplete **(a)**, but CoPA axons became completely ensheathed during the same imaging period (**b)**. Scale bar = 5 μm. **c** Quantification of the percent length of individual axons ensheathed at each time point. t = 0 was defined as the onset of initial wrapping for each subtype within the spinal cord segment imaged (see Methods). Each row of the table contains two dots indicating the subtypes being compared with the corresponding p-value testing for difference in myelination rates of those two subtypes. Error bars represent the mean ± SEM. n (# larvae imaged at 0, 24, 48 hour time points) = 11, 14, 25 (*tbx16*^+^); 15, 12, 6 (*phox2b*^+^); 11, 11, 18 (*pitx2c*^+^); 16, 25, 23 (*isl1[ss]*^+^). P-value from ANCOVA testing for overall difference in myelination rate was 0.0015. Reported p-values for pairwise comparisons were generated with Tukey adjustment.

### Alteration of target axon availability reveals subtype-specific control of initial ensheathment

Previous studies indicate that zebrafish spinal cord oligodendrocytes survive and continue to myelinate in the absence of reticulospinal axons [30]. We recognized this as an opportunity to ask whether the rate that oligodendrocytes myelinate local spinal neuron axons can be influenced by the availability of reticulospinal axons, or is a hard-wired property imposed by the spinal neurons themselves. We first determined the proportion of myelin dedicated toward each axon population. By comparing the amount of myelin associated with individual transgene-labeled axons to the total myelin, we found that oligodendrocytes dedicate 61.3% of their myelinating resources to reticulospinal axons, but only 8.5% to *tbx16*^+^ and *isl1*^+^ local spinal neurons (Fig. 3a). If reticulospinal target axons were removed, we predicted that oligodendrocytes would shift myelinating resources to remaining target axons if initial axon ensheathment is controlled by an oligodendrocyte-intrinsic program. In contrast, if initial axon ensheathment is governed by axon subtype-specific factors, we hypothesized that reducing the abundance of target axons would not alter the initial ensheathment of remaining target axons. In order to alter target axon availability, we took advantage of the different anatomical positions of reticulospinal versus local spinal sensory and interneuron populations. Specifically, because reticulospinal neuronal cell bodies reside in the midbrain and hindbrain, we reasoned that anterior axon ablation near the hindbrain-spinal cord boundary (somites 6-7) would remove distal reticulospinal axon segments in the posterior spinal cord (somites 16 or beyond) (Fig. 3b). We therefore used an anterior spinal cord injury approach to ablate reticulospinal axons in 72 hpf embryos (Additional Fig. 2). To verify that injured axons did not regenerate into posterior segments within 24 hours we directly observed *pitx2c*^+^ reticulospinal axons at 24 hpi. In all instances, we detected either complete axonal loss or remnants of axons consistent with Wallerian degeneration in posterior segments (Fig. 3c, Additional Fig. 2a). In comparison, in control larvae (non-injured siblings), we routinely detected 2-7 *pitx2c*^+^ nMLF axons per spinal hemisegment in posterior segments. In contrast to the complete loss of reticulospinal axons, we observed no changes to the number nor morphology of *isl1[ss]*^+^ RB sensory or *tbx16*^+^ CoPA neurons in posterior segments of injured larvae at 24 hpi (Fig 3c, Additional Fig. 2b). Reticulospinal ablation caused a pronounced effect on animal behavior, because 41/41 animals completely lacking *pitx2c*^+^ nMLF axons in posterior segments also lacked spontaneous swim behavior.

**Fig. 3.**
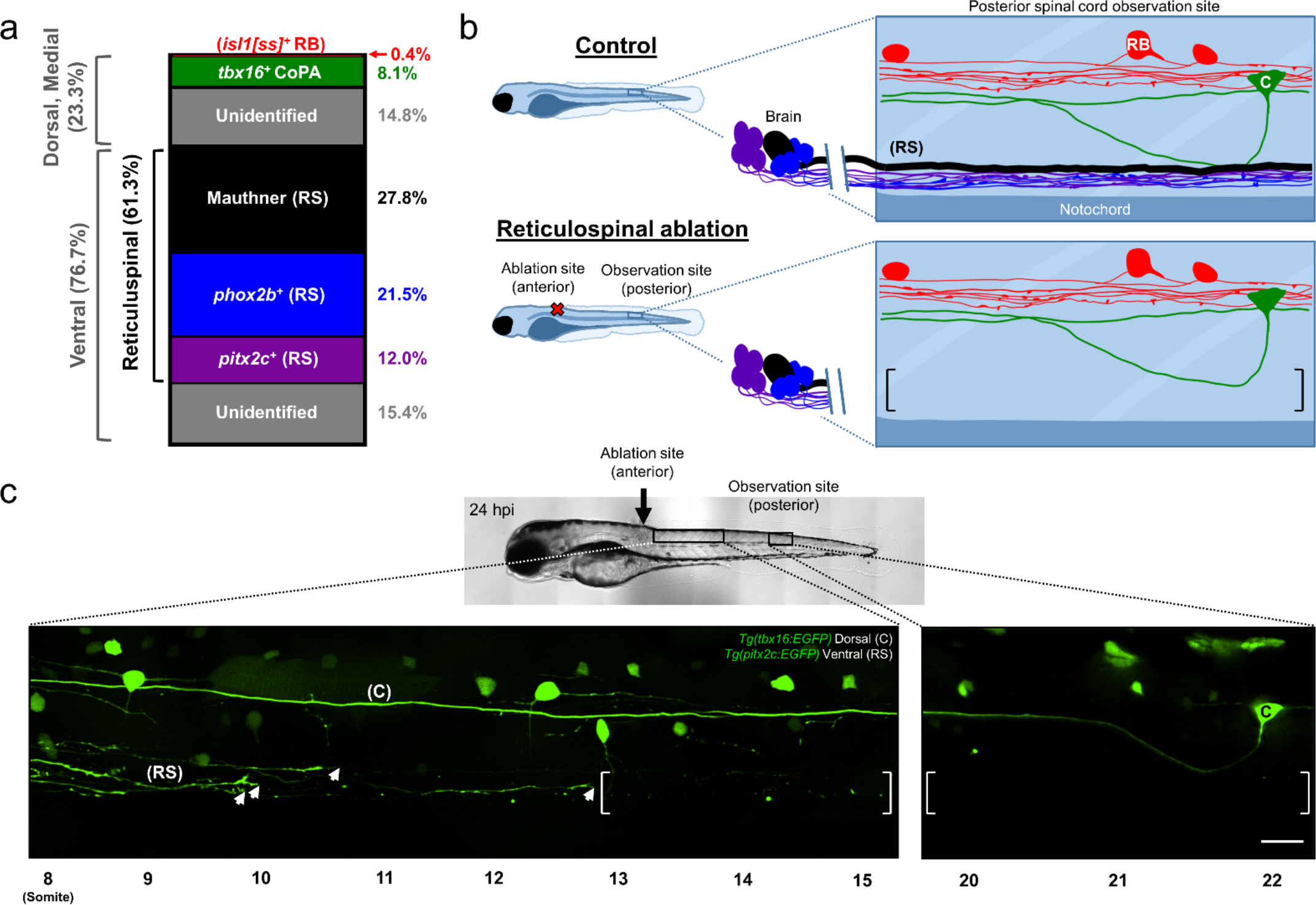
Reticulospinal ablation creates a target axon-deficient environment. **a** Stacked bar graph represents the proportion of myelin dedicated to transgene-labeled subtypes in the posterior spinal cord. n (larvae) = 10 (*isl1[ss]*^+^), 7 (*tbx16*^+^), 13 (Mauthner), 10 (*phox2b*^+^), 8 (*pitx2c*^+^). **b** Schematic depicting reticulospinal ablation. RB, C, and (RS) mark *isl1[ss]*^+^ RB cell bodies, *tbx16*^+^ CoPA cell bodies, and reticulospinal axons, respectively. Brackets enclose an area devoid of reticulospinal axons in ablated animals. **c** Lateral view images of the larval spinal cord acquired at 24 hpi (96 hpf) show severed, regenerating *pitx2c*^+^ axons alongside Wallerian degeneration of their former distal axon segments (left brackets). Arrowheads indicate growth cones of regenerating *pitx2c*^+^ axons, and brackets enclose areas devoid of reticulospinal axons. Note that severed axons never regenerated into posterior observation segments within 24 hours of injury. Therefore, the posterior observation site lacked *pitx2c*^+^ descending reticulospinal axons, but possessed *tbx16*^+^ CoPA ascending local spinal interneurons. Images are tiled confocal maximum z-projections with dorsal up and anterior left. Scale bar = 20 μm.

We next evaluated the validity of this approach to study myelin ensheathment within the posterior spinal cord. Within posterior segments, reticulospinal ablation caused a subtle reduction in *sox10:TagRFP*^+^ oligodendrocyte-lineage cells (somites 24-26, control group = 29 ± 6, ablation group = 23 ± 5 cells per hemisegment, see Additional Fig. 3a,b) and an overall reduction in number of *Tg(sox10:RFP-CaaX)* myelin sheaths (somites 24-25, control group = 34 ± 3, ablation group = 20 ± 3 sheaths per hemisegment, Additional Fig 3d). Notably, we observed a robust loss of sheaths in the ventral spinal cord, where reticulospinal axons normally reside, but no change to sheath numbers in the medial- or dorsal-most spinal cord regions (Additional Fig. 3d). Previous reports using a genetic rather than physical ablation to remove reticulospinal axons from the posterior spinal cord paralleled our observed decrease in oligodendrocyte-lineage cells and myelin sheath numbers [25,30]. We observed reduced average sheath lengths across all spinal cord regions in ablated versus control groups, which could be the result of oligodendrocytes forming more but shorter sheaths on select remaining target axons [25] (Additional Fig. 3e). Taken together, we conclude that ablation creates an environment with a disproportionate reduction in oligodendrocyte-lineage cells (20.7% reduction) compared to target axons (reticulospinal target axons receive 61.3% of myelin). Consequently, axon ablation increased the oligodendrocyte to target axon ratio, and thereby allowed us to test for axon subtype-level alterations in myelination in an environment with excess myelinating potential.

When the oligodendrocyte to target axon ratio is increased, does an oligodendrocyte-intrinsic program continue to maximize surplus myelinating resources by increasing the extent or rate that remaining local spinal neurons become myelinated? We next tested this possibility by interrogating ablation-induced changes to initial ensheathment at the subtype-specific level. Our earlier observations indicated that *isl1[ss]*^+^ RB sensory axons are myelin-competent, but are incompletely myelinated during larval stages (see Fig. 2a). Because overall ensheathment of *isl1[ss]*+ RB axons remained stagnant from 3-5 dpf under baseline conditions (Fig. 2a,c), we reasoned that reticulospinal ablation could shift and accelerate myelination onto RB axons. To test this, we crossed *Tg(isl1[ss]:Gal4, UAS:DsRed)* to the oligodendrocyte reporter *Tg(nkx2.2a:EGFP-CaaX).* At 72 hpf, oligodendrocytes within and posterior to somite 21 had not initiated myelination in the DLF, where RB axons resided (31/31 animals). We therefore performed anterior spinal cord injuries at 72 hpf and observed RB axon ensheathment in all somites within and posterior to somite 21 at 96 hpf. Because ensheathment had not commenced at the time of ablation, all sheaths observed and counted within and posterior to somite 21 had formed after ablation of reticulospinal axons. During the 24-hour period from 72-96 hpf, we observed an average of 2.8 ± 0.5 new sheaths formed onto RB axons in control animals (Fig. 4a,b). Notably, we detected no significant difference in the number nor lengths of new sheaths formed onto RB axons following ablation (Fig. 4b,c).

**Fig. 4.**
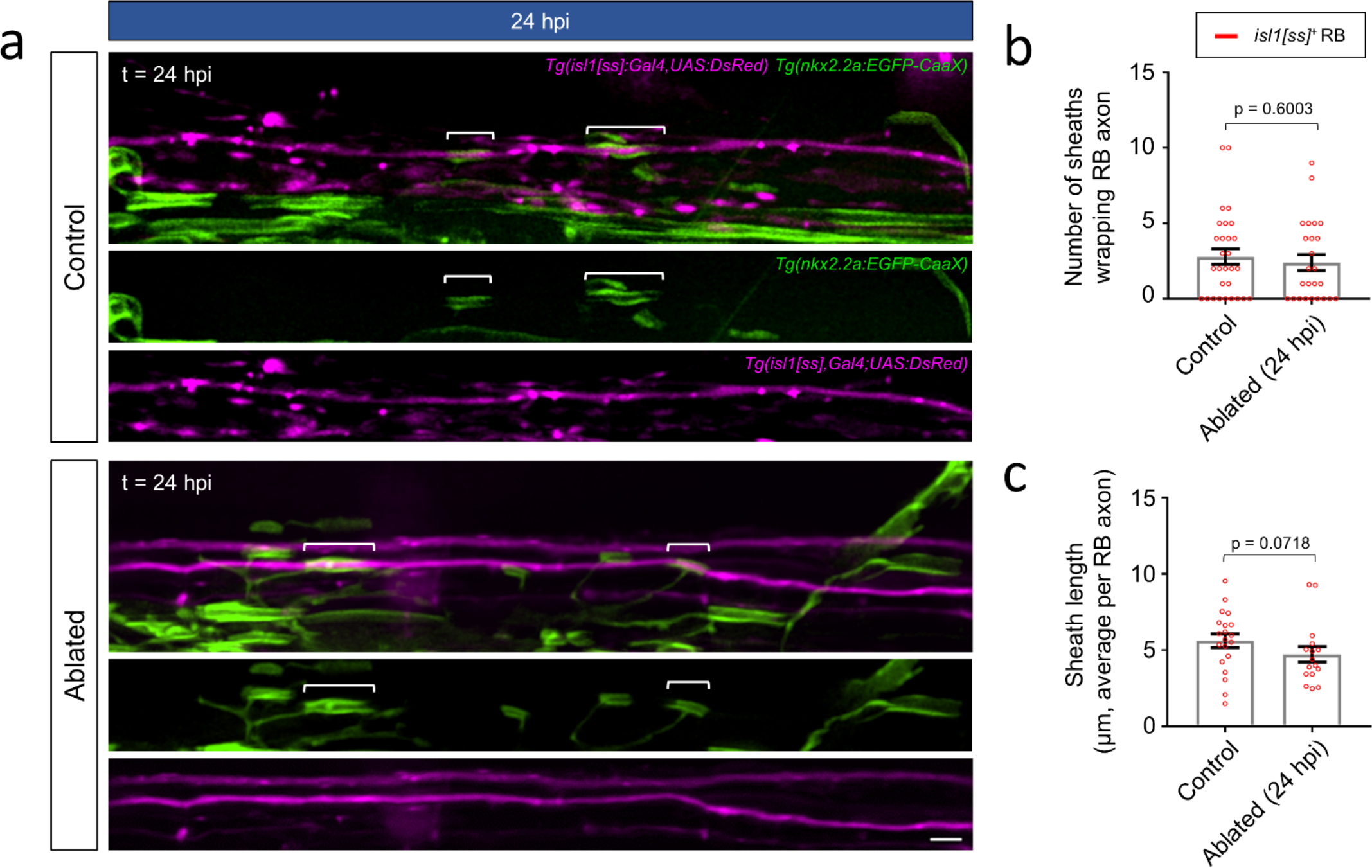
RB spinal sensory neuron axons maintain minimal ensheathment in a target axon-deficient environment. **a** Lateral view images of the posterior spinal cord show ensheathment of RB sensory axons in control (non-injured siblings) and ablated larvae (24 hpi, 96 hpf). Note that oligodendrocytes had not yet formed sheaths within this spinal cord segment at the time of ablation, and therefore all sheaths observed in the ablated group were formed in a reticulospinal axon-devoid environment. Brackets indicate individual sheaths wrapping RB axons. Scale bar = 5 μm. **b** Quantification of RB axon ensheathment. Scatter plot points represent the total number of sheaths wrapping *isl1[ss]:DsRed*^+^ RB axons in the posterior spinal cord of individual larvae (posterior to somite 2). n (larvae) = 29 (control), 25 (ablated). **c** Quantification of average sheath lengths. Scatter plot points represent the average length of all sheaths wrapping RB axons within an individual larvae. n (larvae, total sheath #) = 20, 80 (control) and 16, 60 (ablated). For **b-c**, error bars represent mean ± SEM and p-values report Mann-Whitney test.

To further test the hypothesis that target axon removal would increase the rate of ensheathment of remaining local spinal neuron axons, we next investigated the ensheathment rate of individual *tbx16*^+^ CoPA axons in the presence and absence of descending reticulospinal axons. In contrast to *isl1[ss]*^+^ RB axons, CoPA axons become rapidly ensheathed during the 48-hour period after initial wrapping (Fig. 2b,c), motivating us to assess multiple time points to identify potentially subtle changes to the rate of CoPA axon ensheathment. At both 24 and 36 hpi, CoPA axon myelination was indistinguishable between control and ablated groups (Fig. 5a-d). We observed no effect on the overall percentage length of individual CoPA axons myelinated nor the density of sheaths per individual CoPA axon, together indicating that oligodendrocytes did not respond by dedicating surplus myelin resources toward CoPA axons.

**Fig. 5.**
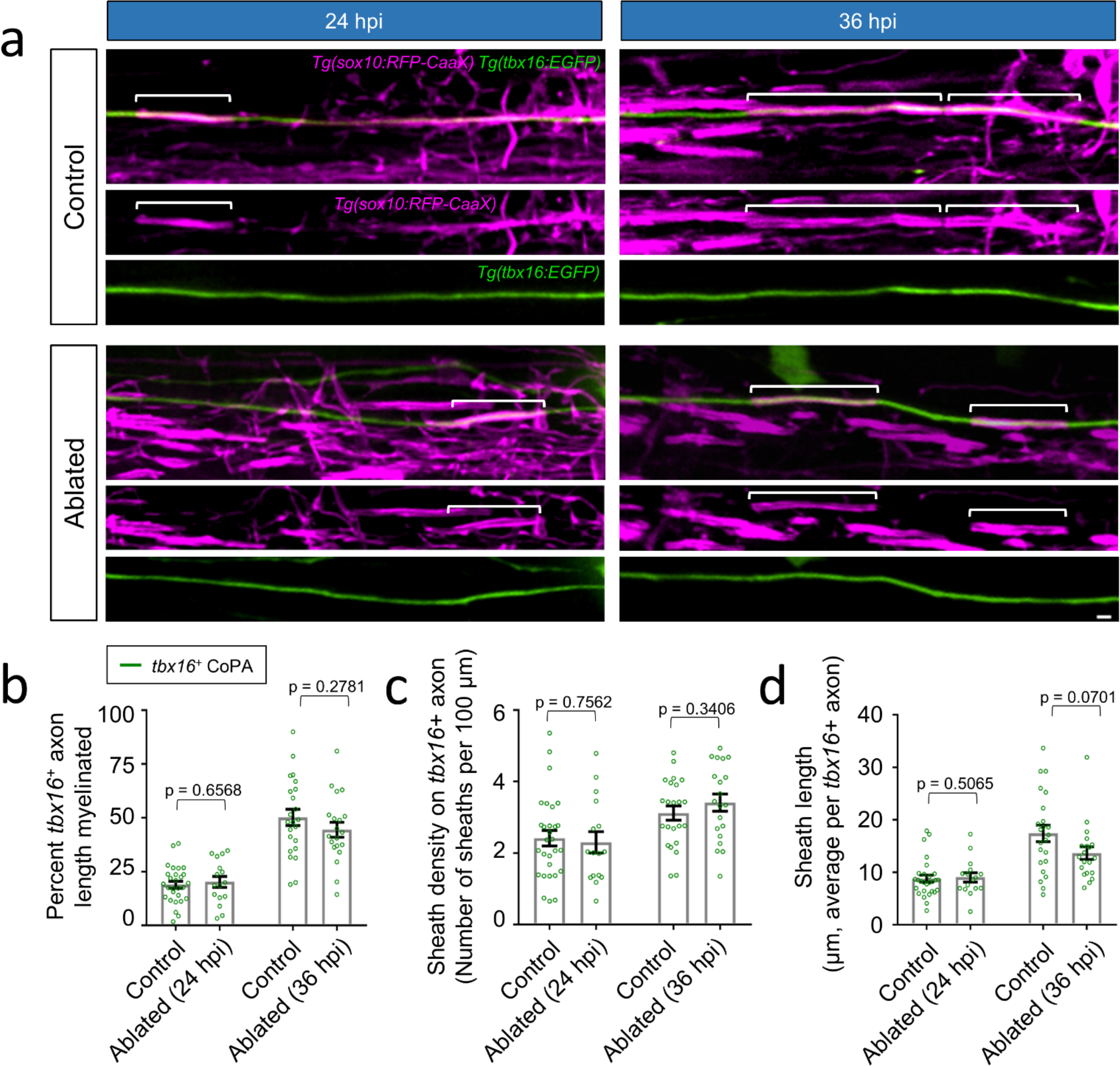
CoPA spinal interneuron axons maintain a constant ensheathment rate in a target axon-deficient environment. **a** Lateral view images of the posterior spinal cord show ensheathment of CoPA axons in control (non-injured siblings) and ablated larvae. The indicated time points (24 and 36 hpi) were acquired at 72 and 84 hpf, respectively, and acquired in a constant spinal cord segment (somites 16-17). Note that ablation was performed prior to the initiation of oligodendrocyte differentiation and axon wrapping (48 hpf), and therefore all sheaths observed and counted formed after ablation of descending reticulospinal axons.
Brackets indicate individual sheaths wrapping CoPA axons. Scale bar = 2 μm. **b-d** Quantification of CoPA axon ensheathment (**b-c**) and overall sheath lengths (**d**). Scatter plot points represent measurements of individual axons **(b-c)** or total number of sheaths wrapping *tbx16*^+^ CoPA axons **(d)** and error bars represent the mean ± SEM. n (larvae, total sheath #) = 28, 94 (control 24 hpi); 16, 53 (ablated 24 hpi); 23, 99 (control 36 hpi); 20, 92 (ablated 36 hpi). Reported p-values were generated using a two-tailed, unpaired t-test to compare groups at matched time points.

Collectively, these findings indicate that when sub-populations of axons are removed, both *tbx16*^+^ CoPA and *isl1[ss]*^+^ RB axons impose a defined rate of ensheathment, and oligodendrocytes do not adaptively shift myelinating resources onto these subtypes.

## Discussion

Oligodendrocytes possess the ability to wrap and form immature myelin without active instruction from axons or synthetic substrates [16–19,31]. This raises the possibility that during normal neural development, oligodendrocytes deploy an intrinsic ensheathment program followed by axon-dependent and adaptive refinement of myelination [20]. In this sequential model, oligodendrocyte processes would indiscriminately wrap segments of any target axon competent for myelination. During this first intrinsic stage, any regional differences in the properties of myelin sheaths, such as sheath lengths, could be explained by the developmental origin and heterogeneity of oligodendrocytes and physical cues such as axon caliber[32]. At a later point during development or adulthood, oligodendrocytes would initiate an adaptive phase whereby axon-derived signals over-ride the oligodendrocyte-intrinsic ensheathment program, and further modify myelin sheath parameters such as length and thickness. Our findings add to a growing body of studies, discussed at length below, which collectively modify or fail to support the sequential model of oligodendrocyte-intrinsic myelination followed by adaptive refinement. If there are two sequential phases, our data indicate that the first phase is not solely mediated by an oligodendrocyte-intrinsic program, but rather is influenced by properties of individual axons. We found that during the initial hours of nascent sheath existence, properties of individual axons differentially influence sheath growth and stabilization, and consequently the rate that individual axon subtypes become myelinated. In this way, the oligodendrocyte-intrinsic ensheathment program and axonal regulators cooperate to determine which axons become myelinated and to what extent. We also show that properties of individual axons are sufficient to prevent changes to their rate of myelination, suggesting that axons autonomously control their own rate of myelination. Taken together, our findings support a cooperative and concurrent model, where from the onset of axon initial wrapping and sheath biogenesis, axon-derived signals override the oligodendrocyte-intrinsic program to direct or restrict the early growth and stabilization of nascent sheaths.

### Axonal influence on oligodendrocyte behavior during initial wrapping and myelination

If myelination is a multi-step process mediated by separable control mechanisms and cell behaviors, when and how do axonal signals participate in CNS myelination? Many neuronal factors influence oligodendrocyte development and myelination, but to date, no single neuronal factor has been identified as necessary to initiate and form myelin sheaths. Although it is possible that such factors will be identified by future studies, an alternative possibility is that oligodendrocytes will initiate wrapping and myelination of any candidate target axon that possesses a supra-threshold axon diameter (> 0.3 μm) and lacks inhibitory signals [17,22], reviewed by [9,13,14]. Our studies do not directly address or exclude this possibility, and it will be important for future studies to determine if the transition from oligodendrocyte-axon contact to initial wrapping is influenced by axonal signals, or is solely mediated by an oligodendrocyte-intrinsic program in vivo. Regardless of whether axon identity influences the initiation of wrapping or subsequent sheath behavior, it is now clear that in both mouse and zebrafish models, individual axon subtypes have acquired distinct myelination patterns very early after the onset of myelination [33].

If the initial growth and stabilization of nascent sheaths is controlled by an oligodendrocyte-intrinsic program, we predicted that nascent sheath behavior would be independent of the axon subtype being myelinated. Instead, we observed that both nascent sheath growth and stabilization depended on the axon subtype being myelinated. For instance, spinal cord commissural axons (*tbx16*^+^ CoPA) supported more rapid sheath growth rates in comparison to sensory (*isl1[ss]*^+^) and reticulospinal axons (*pitx2c*^+^ and *phox2b*^+^). Commissural and sensory axons consistently instructed nascent sheath stabilization, but in comparison, sheaths wrapping reticulospinal axons pruned more frequently. We interpret these observations to mean that axon subtype-specific factors over-ride the oligodendrocyte ensheathment program during the earliest steps of myelination. Because we directly observed the behaviors of nascent sheaths within their initial few hours after formation, if a purely oligodendrocyte-intrinsic program does temporally precede the emergence of axonal influences, our data indicate that these control mechanisms would be separated by only a few hours or less. One alternative explanation could be that the axon subtypes observed are surrounded by distinct extracellular matrix molecules, which could thereby influence the behavior of oligodendrocyte processes and nascent sheaths in an axon-independent manner. Although this is possible, one reason this may be unlikely is that commissural and sensory axons are juxtaposed within the same dorsal spinal cord tract (DLF), yet became myelinated at different rates and to different extents.

Beyond our study, other data further support the interpretation that axonal factors regulate early sheath growth and stabilization. For instance, tetanus-sensitive synaptic vesicle release regulates nascent sheath growth and stabilization, suggesting that axon-derived factors influence early sheath behaviors [5,34,35]. Neuronal activity and transmitter release may be integrated at the level of second messengers within nascent myelin sheaths, because specific Ca2+ patterns associate with sheath growth or retraction [36,37]. Notably, in these studies as well as our own work reported here, sheath behaviors were observed at early developmental stages immediately following initial axon wrapping. Axon-dependent regulation of sheath behavior would not be expected to occur at this stage if initial myelination were solely controlled by an oligodendrocyte-intrinsic program, and adaptive or axon-dependent control would instead emerge at a later developmental time point. Such activity-dependent and synaptic vesicle regulation at the early stages of sheath formation and growth suggest that experiences and adaptive changes to myelin can occur from the onset of myelination, calling question to the notion of separable and sequential hardwired then adaptive phases of myelination.

The subtype-specific differences in nascent sheath growth rate and pruning frequencies we observed may explain how some axons become myelinated more rapidly than others. For instance, the rapid growth rates and absence of sheath retractions associated with spinal cord commissural axons would predict rapid overall ensheathment, which is consistent with our observation that *tbx16*^+^ CoPA axons became completely myelinated within the 48 hours after initial wrapping. Conversely, the comparatively slower sheath growth rates and increased pruning frequencies among nascent sheaths wrapping reticulospinal axons corresponded to a reduced overall rate of myelination during the same time period. Because spinal cord commissural and reticulospinal axons possess indistinguishable diameters at this developmental stage [23], these differential sheath behaviors are most likely not attributable to differences in axon caliber, and instead support the notion that axon-derived molecular signals are active regulators of sheath behavior during the earliest phases of myelination. These behaviors may explain the emergence of unique myelination patterns on distinct neuronal subtypes, such as those observed on axons positioned adjacent to one another within the mouse neocortex [33].

### Can axons autonomously control their own myelination?

The observed differences in the rates that individual axon subtypes become myelinated could be accomplished by one or more control mechanisms. First, if all myelin-competent axons are comparable targets for oligodendrocytes, then the rate that any individual axon becomes myelinated could be based on the local supply of oligodendrocytes and their limited myelinating capacity. Pointing against this possibility, we found that commissural and sensory axons become myelinated at vastly different rates, despite the fact that these axons both reside within the DLF. Furthermore, in the mouse cortex, myelination of distinct axon types initiates at different timepoints which would not be expected if myelination were solely determined by local supply of oligodendrocytes [33]. As an alternative to the oligodendrocyte supply model, the molecular properties of an individual axon could serve to define or instruct its own rate of myelination. In support, Auer et al. reported that following oligodendrocyte ablation, regenerated myelin tends to restore the same pattern and position of myelin that was found on individual axons prior to oligodendrocyte death [8]. Our axon ablation experiments also contribute to this question. We reasoned that if axons lacked the ability to dictate their own set point rate of myelination, they would become more rapidly and fully myelinated when the oligodendrocyte to target axon ratio was increased. Instead, we found that both spinal commissural and sensory axons maintained a constant rate of myelination, despite the surplus of oligodendrocytes relative to remaining target axons. Almeida et al. performed similar experiments but used a genetic approach rather than physical ablation to reduce the number of reticulospinal target axons within the posterior spinal cord. In this study oligodendrocytes did not increase myelination of RB sensory axons, but did occasionally perform ectopic myelination of cell bodies [25]. Inverse experiments that increased the target axon supply caused oligodendrocytes to increase myelinating potential in order to meet the increased demand imposed by supernumerary target axons, and ensure the normal rate of myelination for this reticulospinal (Mauthner axon) subtype [38]. Taken together, these studies indicate that some neuronal subtypes are hard-wired to instruct oligodendrocyte behavior during the initial stages of myelination, and further suggest that the mechanisms preventing ectopic or accelerated axon ensheathment may be dominant to those preventing myelination of cell bodies and dendrites.

### Implications for myelin plasticity and regeneration

Oligodendrocytes and their progenitors detect and respond to changes in the local environment in various ways to ensure the intended distribution of myelin onto target axons. As recently discussed, increasing the number of target axons causes individual oligodendrocytes to increase their myelinating potential, but may not influence specification, proliferation, or differentiation of oligodendrocyte progenitor cells (OPCs) and oligodendrocytes [38]. In contrast, when the number of target axons is reduced, oligodendrocytes continue to differentiate but can be reduced in number due to apparent axon-dependent influences on OPC proliferation and survival [30,39]. How oligodendrocytes sense the number of target axons and relay this to the mitotic and apoptotic pathways is incompletely understood, but these mechanisms may exist to ensure that the proper amounts of myelin are formed at the right times and places during development, and may also enable adaptation to neuronal changes.

Our studies of how ensheathment of individual axons is influenced by changes to the availability of other target axons also has implications for myelin plasticity. Are the early phases of myelination mediated by hard-wired developmental processes that are controlled exclusively by gene regulatory networks intrinsic to neurons and oligodendrocytes? Alternatively, could experience-based changes modify gene regulatory networks and pathways controlling initial sheath growth, stabilization, and the extent and rate that individual axons become myelinated? These questions are incompletely resolved, but accumulating evidence indicate that multiple forms of distinct adaptions can occur, and that such changes may depend on developmental stage, anatomical region, and axon subtype identity [6,23,40,41]. Differences in neuronal excitability between neighboring axons may influence the extent that individual axons become myelinated [5,23,35,42]. Our observations that spinal commissural and sensory axons restrict additional myelin sheaths or more rapid ensheathment, even when myelinating potential is in excess, suggest that these subtypes would not invite or accept additional myelin if activity were altered in neighboring axons. Instead, our findings are consistent with the notion that any adaptive shifts in-between axon subtypes occurring in response to altered activity or experience are likely to be highly restricted to specific subtypes, rather than widespread across all subtypes. For example, reticulospinal axons may permit surplus oligodendrocytes to form more, albeit shorter myelin sheaths along their length in an adaptive manner [25]. We did not observe this effect on CoPA and RB sensory axons in our study. Notably, this correlates with the participation of synaptic vesicle release in the myelination of reticulospinal axons, but not for CoPA axons [23].

If axons provide critical information that instructs early myelin sheath growth and stabilization, this suggests that strategies and regenerative therapies focused solely on manipulating oligodendrocyte biology may be inefficient or insufficient to induce remyelination. Instead, if the properties of demyelinated axons have changed, these axonal alterations may represent important targets in addition to promoting oligodendrocyte differentiation and myelinating potential. For example, emerging evidence indicates that axonal galectins mark non-myelinated axon segments, and that galectins may be mis-expressed within Multiple Sclerosis lesions [43,44]. Numerous additional negative regulators of myelination have been identified that if targeted, could mitigate the barriers to remyelination (reviewed by [9,13,14]). It will be interesting for future studies to identify spinal cord commissural axon properties responsible for the superior sheath stabilizing and growth inducing properties of this subtype in comparison to reticulospinal and spinal sensory axons. One possibility is that the levels of second messenger and kinase cascades within axons could account for such differences. For example, increased activation of Akt-mTOR signaling within cerebellar parallel fiber axons is sufficient to cause ectopic myelination of this normally unmyelinated population [21]. Because activating axonal pathways such as the Akt-mTOR pathway is also sufficient to increase axon diameter, such manipulations may be sufficient to induce both permissive and instructive axonal cues, and therefore may represent promising avenues for myelin regeneration.

## List of abbreviations

CNS: Central nervous system
CoPA: Commissural primary ascending
DLF: Dorsal longitudinal fasciculus
dpf: Days post-fertilization
hpf: Hours post-fertilization
hpi: Hours post-injury
nMLF: Nucleus of the medial longitudinal fasciculus
OPC: Oligodendrocyte progenitor cell
RB: Rohon-Beard

## Acknowledgements

We thank Mary Halloran (University of Wisconsin) for the *Tg(pitx2c:EGFP)* line, Michael Lardelli (University of Adelaide) for the *Tg(tbx16:EGFP)* line, and Angie Ribera (University of Colorado – Anschutz Medical Campus) for the *Tg(isl1[ss]:Gal4-VP16, UAS:DsRed)* line. We also thank John Henley and Craig Nelson (Mayo Clinic) for spinal cord transection assistance, Erika Vail and Mary Diekmann for technical assistance, and Kevin Okome and Nina Horabik for statistical consulting.

## Author Contributions

J.H.H., H.N.N., A.J.T., M.R.M., and A.J.K. designed research, all authors acquired and analyzed data, J.H.H. and H.N.N. wrote the manuscript, and all authors edited the manuscript.

## Availability of data and materials

All data, zebrafish lines, plasmids, and other reagents are available upon request from the corresponding author.

## Methods

### Zebrafish lines and husbandry

All animal work performed in this study was approved by the Institutional Animal Care and Use Committee at Winona State University. Zebrafish embryos were raised at 28.5° C in egg water (0.0623 g/L Coralife marine salt) and staged according to hours post-fertilization or morphological criteria. The sex of animals was not determined. Transgenic lines used in this study included *Tg(sox10:RFP-CaaX)^vu234^*, *Tg(sox10:TagRFP-T)^co26^*, *Tg(nkx2.2a:EGFP-CaaX)^vu16^*, *Tg(phox2b:EGFP) ^w37^*, *Tg(phox2b:GAL4) ^co21^*, *Tg(tbx16:EGFP) ^uaa6/812c^*, *Tg(isl1[ss]:Gal4-VP16, UAS:DsRed) ^zf234^*, *and Tg(pitx2c:EGFP) ^zy8^*.

### Microscopy and quantitative image analysis

Embryos or larvae were embedded in 1% low-melt agarose using tricaine (MS-222) anesthesia (3.48 mM, Pentair). All images were acquired or rotated to show lateral views with anterior left and dorsal up. Confocal microscopy was performed using an Olympus IX-81 equipped with a disk spinning unit (DSU), 60× 1.3 NA silicone immersion objective (unless otherwise stated), LED illumination with narrow bandpass filters, and a Hamamatsu Orca-R2 CCD camera. Z-intervals were 400 nm and individual camera pixel sizes were 107.5 nm, or if 2×2 binning was required for fluorescence detection, 215 nm. All image analysis was performed using CellSens (Olympus) and NIH ImageJ software. Images were deconvolved using 3D iterative deconvolution (CellSens, Olympus) and all projection images were produced using maximum intensity.

### Determination of axon subtype-specific ensheathment, sheath growth rates, and pruning

To determine overall ensheathment rates, a baseline time point was first defined for each individual axon subtype as a specific spinal cord segment where axon wrapping had initiated in the majority of larvae imaged. This baseline (t = 0) was determined by the presence of a low but consistent level of sheath initiation on transgene-labeled axons. Subsequent images were acquired 24 and 48 hours later within the same spinal cord segment. For each image, the percent of individual axon length ensheathed was determined by dividing the sum of all sheath lengths (with GFP/RFP colocalization) by the total axon length within a field of view. For measurements of individual sheath growth and pruning, we collected images of the same nascent sheath at t = 0 and t = 3 hours. A nascent sheath was defined as a structure with processes wrapped around the entire circumference of an axon in x-y and orthogonal views, containing a clearly defined lumen devoid of oligodendrocyte-channel fluorescence, and myelinating internodes 2-5 μm in length. During the three-hour period between image acquisitions, fish were incubated at 28.5°C while embedded in 1% low-melt agarose under tricaine anesthesia. Sheath growth rate measurements were restricted to sheaths that were present at both time points.

### Determination of myelin sheath distribution on individual axon subtypes during early myelination

We first measured the percent of all myelin associated with the Mauthner axon, dorsal to the Mauthner, and ventral to the Mauthner axon. To accomplish this, the combined sheath lengths within the three aforementioned dorsal-ventral domains were summed and then divided by the combined lengths of all sheaths within the spinal hemisegment. Images were acquired in somites 24-25 (restricted to larvae possessing 10-50 sheaths per domain) at 96 hpf (hours post-fertilization).

Individual sheaths within each of the three domains were next assigned to individual axon subtypes. First, axon subtypes were assigned a domain of residence based upon their anatomical position (e.g. *isl1*^+^ RB is dorsal/medial). Next, we determined the lengths of all sheaths within the resident domain. We separated these values into two categories (yes/no) based on whether the sheath exhibited RFP/GFP colocalization with an axon marked by the subtype-specific reporter line. The total length of sheaths with RFP/GFP colocalization was then divided by the combined length of all sheaths within the resident domain. Images were acquired at 98-105 hpf. Images of ventral axon subtypes were acquired at somite 27, whereas images of dorsal/medial subtypes were acquired at somite 21 to account for the slight developmental delay of myelination within the dorsal spinal cord.

### Ablation procedures, validation, and imaging experiments

A sharp-edged ablation needle was generated from pulled capillary glass and adjusted to 60 μm diameter using #5 forceps. Zebrafish embryos staged at 2-3 days post-fertilization (dpf) under tricaine anesthesia were transferred atop Sylgard-coated petri dishes and aligned anterior left and dorsal up, with minimal egg water, under a dissecting microscope. The ablation needle was held roughly vertical and perpendicular to the embryo (60-75 degree angle) and inserted through the spinal cord at somite 7. Care was taken to avoid desiccation of the animal or injury to the notochord. Animals were immediately transferred to egg water and incubated at 28.5°C. Behavior was assessed at 24 hours post-injury (hpi) and immediately prior to data acquisition. Animals exhibiting any signs of spontaneous swim in a period of 20 minutes, response to light stimulation, or response to touch (pin tool) near the otolith were discarded. Unless otherwise indicated, the *Tg(pitx2c:EGFP)* reporter was crossed into the background of all animals used for experiments to visually verify the absence of distal reticulospinal axon segments in the region of interest prior to data acquisition. Because the absence of spontaneous swim behavior and complete morphological reticulospinal axon ablation corresponded (41 of 41 animals), behavioral criteria were used to select animals for imaging of *isl1[ss]*^+^ Rohon-Beard (RB) axons because *nkx2.2a:EGFP-CaaX*^+^ cells prevented observation of GFP^+^ reticulospinal axons in the ventral spinal cord. Non-injured, stage-matched siblings were used as controls.

Oligodendrocyte-lineage cell counts and myelin sheath counts were performed at 96 hpf (corresponding to 24 hpi), and images were acquired within somites 24-26 and 24-25, respectively. For oligodendrocyte-lineage cell counts, images were acquired using a 20× 0.7 NA objective. The domains were delineated by sheaths wrapping the *tbx16*^+^ CoPA axon, whose diagonal ascending portion traversed the medial domain then projected horizontally along the dorsal domain, and by sheaths wrapping the Mauthner axon, which separated the medial domain (sheaths above) from the ventral domain (sheaths below). Sheaths wrapping the Mauthner axon were excluded from analysis. Local spinal neuron cell body and axon counts were performed at 96 hpf (24 hpi) and images were acquired at or posterior to somite 20 for *isl1[ss]*^+^ neurons and within somites 16-17 for *tbx16*^+^ neurons.

For assessment of *isl1[ss]*^+^ RB sensory axon ensheathment, larvae were imaged at 96 hpf (24 hpi). All sheaths at or posterior to somite 21 were counted and imaged. Selection of somite 21 was based on the absence of sheath initiation at this segment at the time of injury (72 hpf), enabling later assessment of sheaths that had formed in an environment lacking descending reticulospinal axons. For assessment of *tbx16*^+^ CoPA axon ensheathment, the distribution of *tbx16*^+^ CoPA neurons necessitated alterations in somite of data acquisition. Because the *Tg(tbx16:EGFP)* reporter line marks relatively few CoPA axons in posterior spinal cord somites 24-26 (see Additional Fig. 1a), we shifted investigation anteriorly to somites 16-17, where CoPA axons were more abundant. The anterior shift in somite acquisition required a concomitant shift in time points to earlier developmental stages (injury was performed at 48 hpf as opposed to 72 hpf in *isl1[ss]*^+^ RB experiments). Larvae were imaged at 72 hpf and 84 hpf (24 hpi and 36 hpi). To verify injury performed at the earlier 48 hpf time point still resulted in the complete absence of reticulospinal axons within the posterior spinal cord at both 24 and 36 hpi, the *Tg(pitx2c:EGFP)* reporter line was crossed into all animals for internal validation. Zebrafish embryos have not initiated myelination by 48 hpf, therefore any myelin that we observed had formed in the absence of reticulospinal axons.

### Quantification and statistical analysis

In instances where the data were normally distributed we used an unpaired, two-tailed t-test for hypotheses concerning differences between two means. In instances where data were not normally distributed, we preferentially utilized non-parametric tests to compare distributions: the Mann-Whitney test when comparing distributions of two groups or the Kruskal-Wallis test for comparing distributions of more than two groups. When the Kruskal-Wallis test was significant, Dunn’s method was used to adjust for multiple comparisons to determine where differences existed. A chi-square test was used to determine if individual axon subtypes supported sheath pruning versus stabilization (binary outcomes) by comparing percentages of outcomes across groups. Tukey-adjusted pairwise comparisons were then used to determine where differences existed. In order to determine if myelination rate differed across the axon subtypes, an analysis of covariance (ANCOVA) was fit, allowing for an interaction between time and axon subtype. Tukey-adjusted pairwise comparisons were used to assess potential myelination rate differences between individual subtypes and generate p-values. All statistical analyses were performed using Prism 6 (GraphPad) or JMP Pro 14.0. In all figures, error bars represent mean ± standard error of the mean (SEM).

