## Supplemental Figures for "Individual neuronal subtypes control initial myelin sheath growth and stabilization"

### Additional Figures

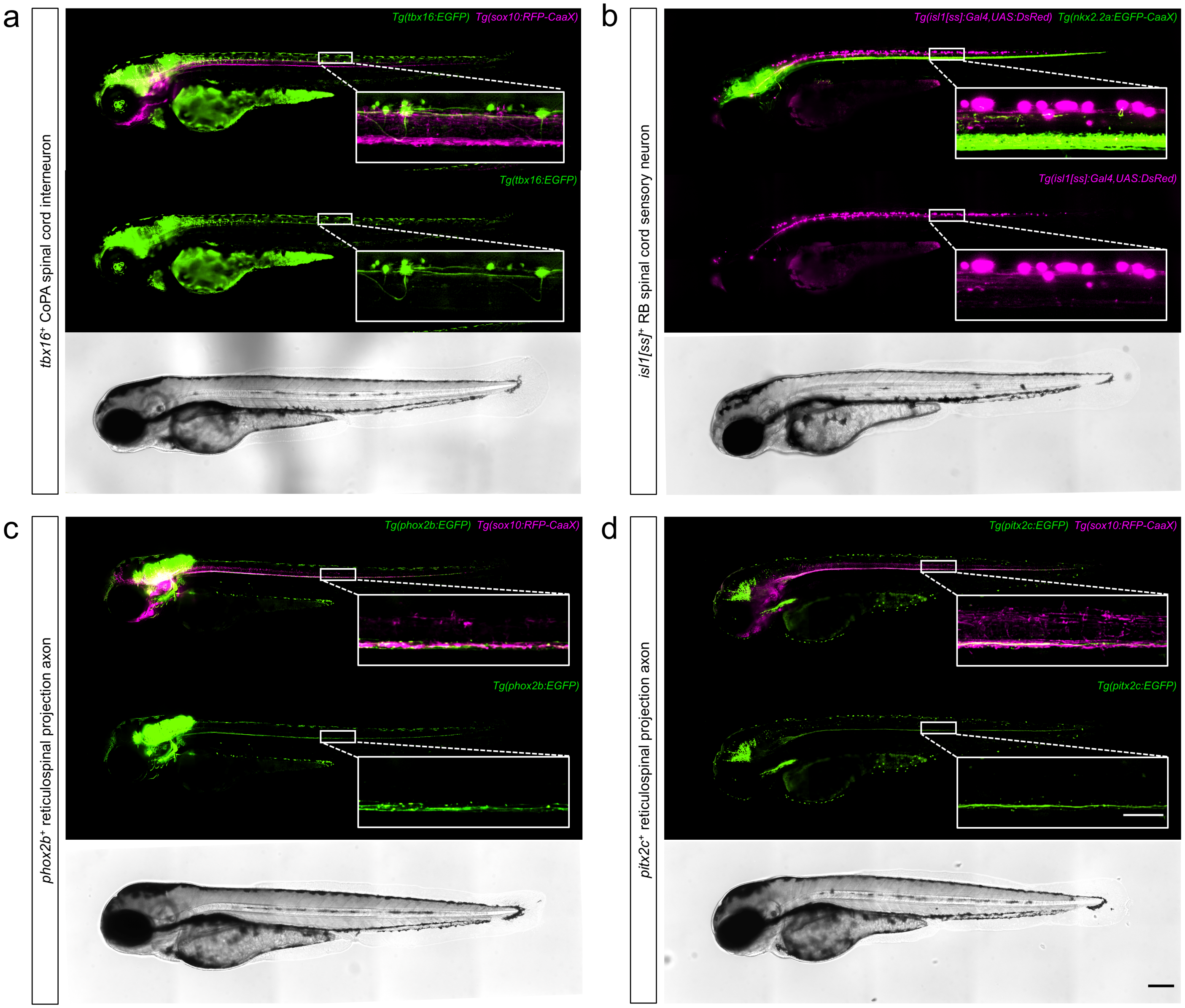


*Additional Fig. 1. Characterization of transgenic reporter lines used.* **a-d** Lateral view images of the larval spinal cord show an overview of transgenic reporters (indicated) used to mark defined axon subtypes or oligodendrocyte membranes. Note that *tbx16*$^{+}$ CoPA and *isl1[ss]*$^{+}$ RB axons project from spinal neurons, whereas the cell bodies of *pitx2c*$^{+}$ and *phox2b*$^{+}$ reticulospinal neurons are located in the brain. Images are tiled confocal maximum z-projections with dorsal up and anterior left. Scale bars = 200 µm (full body) and 50 µm (inset).

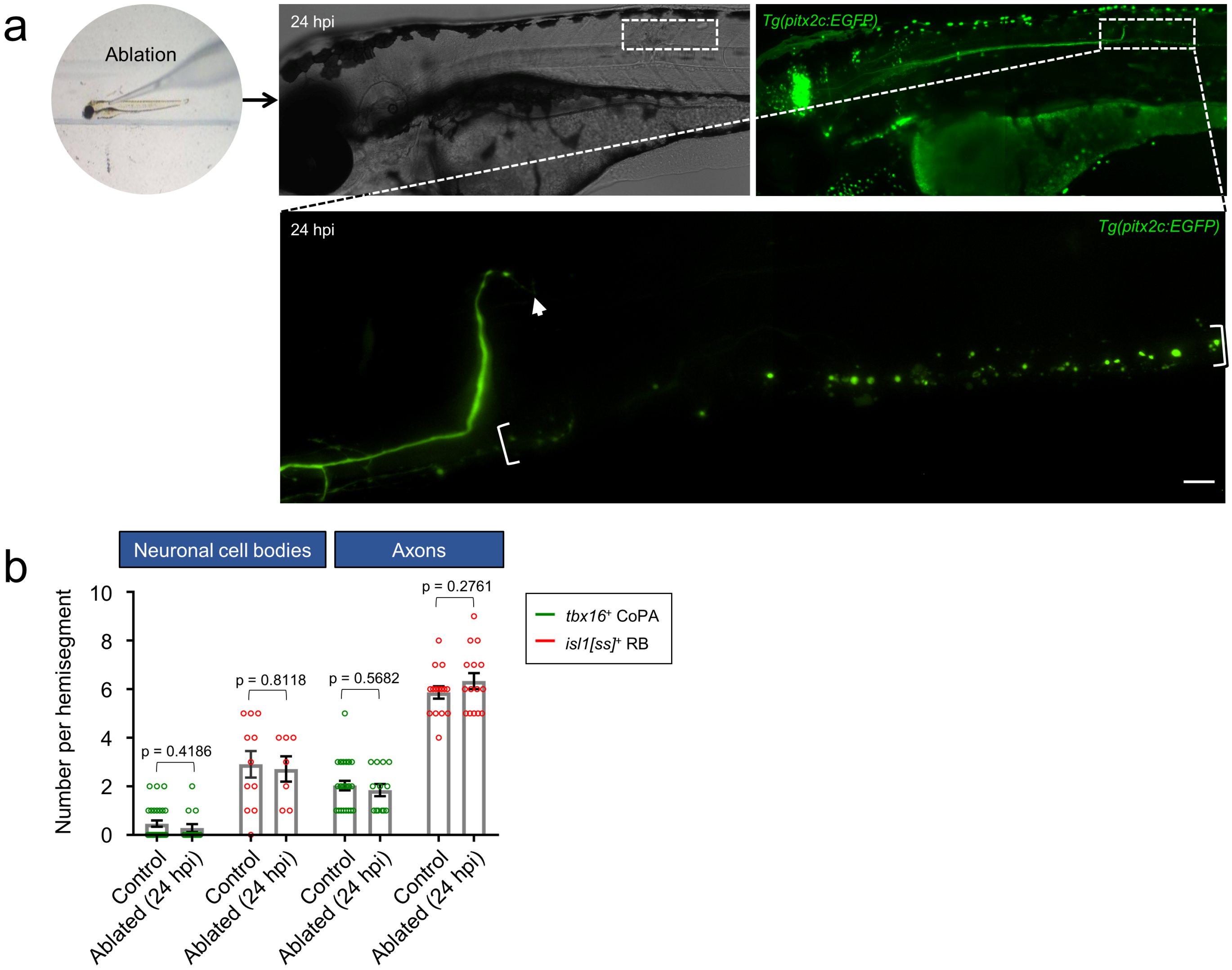


*Additional Fig. 2. Ablation removes posterior reticulospinal axon segments without altering the number of CoPA and RB local spinal cord axons.* **a** Anterior injury (somites 6-7) performed on a 72 hpf larva with a 60 µm glass capillary needle. Lateral view images of the larval spinal cord acquired at 24 hpi (96 hpf) show severed *pitx2c*$^+$ descending reticulospinal axons and Wallerian degeneration distal to the ablation site (bracketed). Images are tiled confocal maximum z-projections with dorsal up and anterior left. Scale bar = 10 µm. **b** Quantification of *tbx16*$^+$ CoPA and *isl1[ss]*$^+$ RB neuron cell bodies and axons in control (non-injured siblings) and ablated (24 hpi, 96 hpf) larvae. For neuron cell body counts, n (larvae) = 28, 14 (*tbx16*$^+$ control, ablated); 11, 6 (*isl1[ss]*$^+$ control, ablated) and for neuron axon counts, n (larvae) = 27, 13 (*tbx16*$^+$ control, ablated); 15, 15 (*isl1[ss]*$^+$ control, ablated). Error bars represent the mean ± SEM. Reported p-values were generated using a two-tailed, unpaired t-test to compare groups at matched time points.

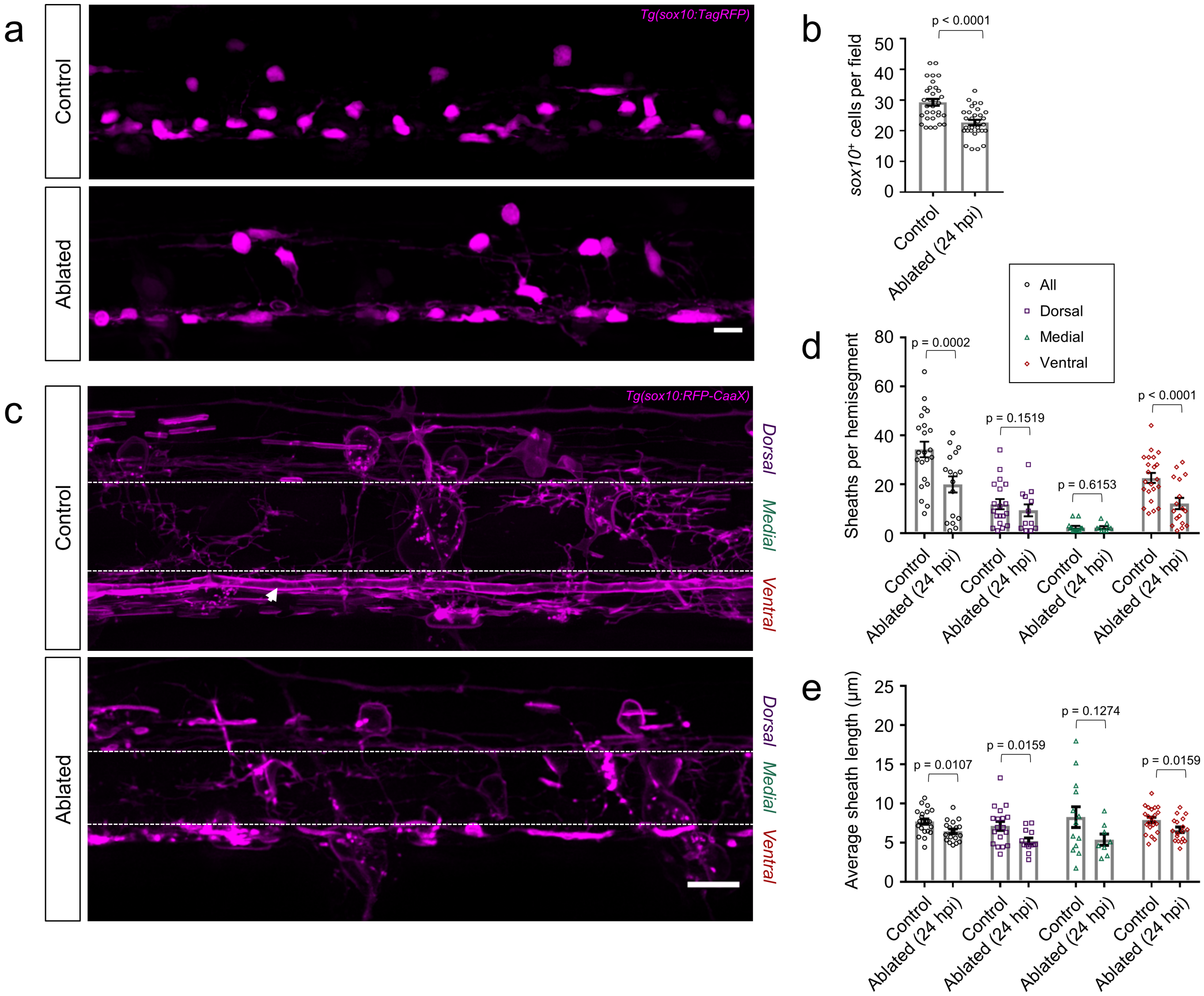


*Additional Fig. 3. Effects of reticulospinal ablation on posterior spinal cord glia.* **a** Lateral view images of the posterior spinal cord show oligodendrocyte-lineage cell bodies marked by *Tg(sox10:TagRFP)* in control (non-injured siblings) and ablated larvae (24 hpi, 96 hpf). **b** Quantification of oligodendrocyte-lineage cell bodies shows the number of *sox10:TagRFP*$^{+}$ oligodendrocyte-lineage cells per field (spinal cord hemisegment). n (larvae) = 32 (control); 31 (ablated). **c** Lateral view images of the posterior spinal cord show myelin sheaths marked by *Tg(sox10:RFP-CaaX)* in control and ablated larvae (24 hpi, 96 hpf). Dashed white lines demarcate dorsal, medial, and ventral domains (indicated). **d-e** Quantification of myelin sheaths **(d)** and average myelin sheath lengths **(e)**. Scatter plot points represent the number of *sox10:RFP-CaaX*$^{+}$ myelin sheaths per spinal cord hemisegment **(d)** and average sheath length per animal **(e)** reported as a total and by domain (dorsal, medial, and ventral). For **d-e**, sheaths wrapping the Mauthner axon (Supplemental Fig. 3c arrowhead) were excluded from data. For sheath counts **(d)**, n (larvae) = 22, 17 (total control, total ablated); 19, 12 (dorsal control, dorsal ablated); 13, 9 (medial control, medial ablated); 22, 17 (ventral control, ventral ablated) and for average sheath lengths **(e)**, n (larvae) = 22, 19 (total control, total ablated); 19, 14 (dorsal control, dorsal ablated); 13, 9 (medial control, medial ablated); 22, 19 (ventral control, ventral ablated). All images are tiled confocal maximum z-projections with dorsal up and anterior left. Scale bars = 10 µm. For all graphs and statistical comparisons, error bars represent the mean ± SEM. Reported p-values were generated using a two-tailed, unpaired t-test to compare groups at matched time points.
